# The flying insect thoracic cuticle is heterogenous in structure and in thickness-dependent modulus gradation

**DOI:** 10.1101/2021.06.30.450643

**Authors:** Cailin Casey, Claire Yager, Mark Jankauski, Chelsea Heveran

## Abstract

The thorax is a specialized structure central to an insect’s ability to fly. In the thorax, flight muscles are surrounded by a thin layer of cuticle. The structure, composition, and material properties of this chitinous structure may influence the efficiency of the thorax in flight. However, these properties, as well as their variation throughout anatomical regions of the thorax or between insect taxa, are not known. In this work, we provide a multi-faceted assessment of thorax cuticle for fliers with asynchronous (honey bee; *Apis mellifera)* and synchronous (hawkmoth; *Manduca sexta*) muscles. We investigated cuticle structure using histology, material composition through confocal laser scanning microscopy, and modulus gradation with nanoindentation. Our results suggest that cuticle properties of the thorax are highly dependent on anatomical region and species. Modulus gradation, but not mean modulus, differed between the two types of fliers. In some regions, *A. mellifera* had a positive linear modulus gradient from cuticle interior to exterior of about 2 GPa. In *M. sexta*, the modulus gradients were variable and were not well represented by linear fits with respect to cuticle thickness. We utilized finite element modeling to assess how measured modulus gradients influenced maximum stress in cuticle. Stress was reduced when cuticle with a linear gradient was compressed from the high modulus side. These results support the protective role of the *A. mellifera* thorax cuticle. Our multi-faceted assessment advances our understanding of thorax cuticle structural and material heterogeneity and the potential benefit of material gradation to flying insects.

**Graphical Abstract:** 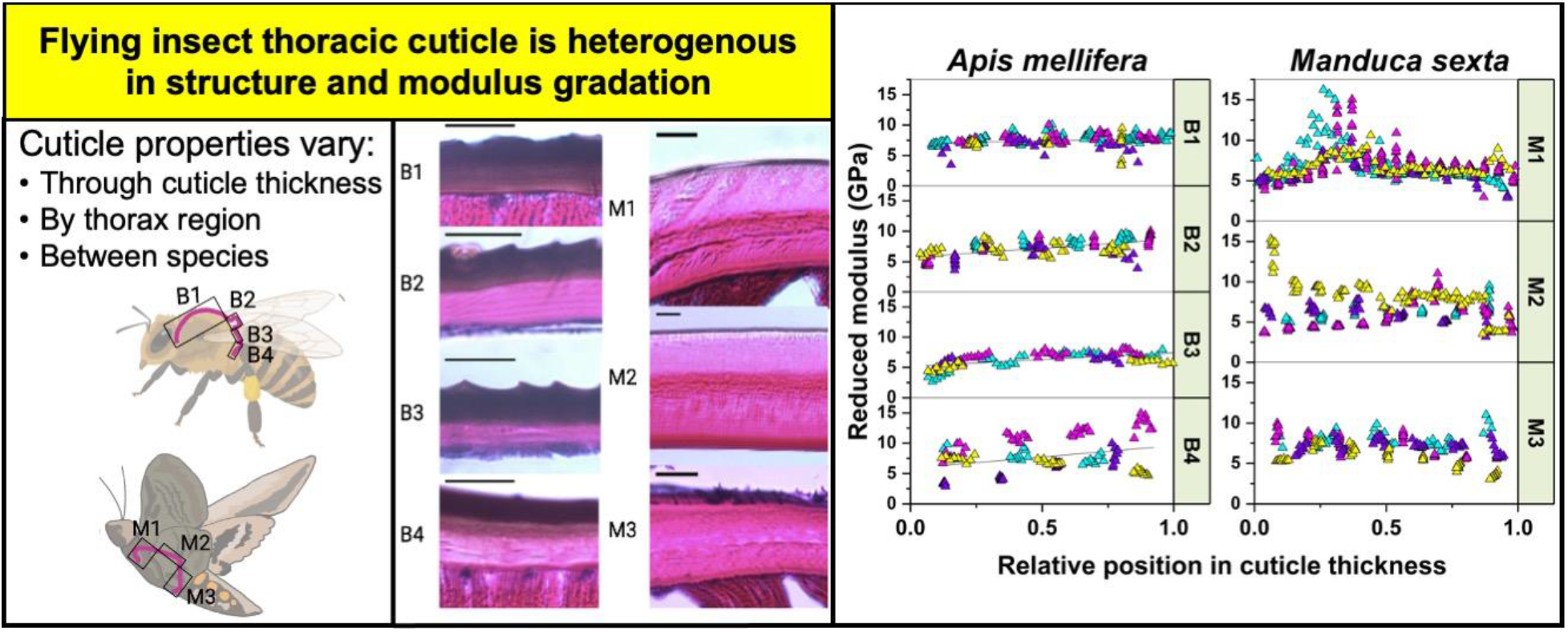

**Statement of Significance:** The insect thorax is essential for efficient flight but questions remain about the contribution of exoskeletal cuticle. We assessed the thorax cuticle using a high resolution multi-faceted approach to determine how cuticle properties vary within thorax anatomical regions and between fliers with asynchronous (honey bee; *Apis mellifera*) and synchronous (hawkmoth; *Manduca sexta*) muscles. We examined structure using histological staining, modulus using nanoindentation, and material composition using confocal scanning light microscopy. We further utilized finite element modeling to understand the effect of the modulus gradations observed experimentally on stress accumulation. Cuticle properties vary through cuticle thickness, by thorax region, and between flight lineages.

## 1. Introduction

As the only invertebrates to evolve flight, insects interest researchers across disciplines. As a group, flying insects can hover, travel long distances, and reach speeds of 25 kilometers per hour [1]. Insects achieve efficiency, especially necessary for hovering, through a mechanism called indirect actuation. In indirect actuation, flight muscles deform the thorax to indirectly flap the wings [2]. The thorax has two main components- flight muscles, and the thin-walled ellipsoidal or box-like exoskeletal cuticle where flight muscles attach [2]. Cuticle is a composite of chitin, water, and proteins which is organized into distinct layers, including endocuticle, exocuticle, and epicuticle [3].

The thorax contains two main flight muscle groups, the dorsal ventral muscles (DVM) and the dorsal lateral muscles (DLM). Flight muscle forces are translated into flapping via intricate connections at the wing hinges [3,4]. Within the indirect lineage, fliers have either *synchronous* or *asynchronous* flight muscles. For the purpose of this paper we will distinguish these insects as asynchronous and synchronous fliers. Synchronous flight requires a neural impulse for each wing beat and is usually associated with flapping rates of <100 Hz [5]. Asynchronous flight produces work in the thorax through delayed stretch activation leading to multiple flaps per impulse [2].

While the physiology of flight muscles has been studied extensively, the cuticle has not. In particular, the roles of thorax cuticle geometry, layering, and material gradation in flight are not well understood. Prior nanoindentation studies found increasing elastic modulus and hardness from the cuticle interior to exterior in the gula [6–8], infrared sensillum [7], elytra [9], and tarsal setae [10] of various beetles, the grasshopper mandible [11], and the locust tibia [12] and sternum [13]. These gradations have been shown to provide protection and improve resilience [14,15]. A modulus gradation in the thorax would be expected to confer the same benefits, though to our knowledge, modulus gradations have not been studied in the thorax cuticle of either synchronous or asynchronous fliers.

The goal of the present work is to assess the thorax cuticle layer organization and modulus gradation for flying insects. We used histology, nanoindentation, and confocal laser scanning microscopy (CLSM) to investigate cuticle layer organization, composition (i.e., presence of resilin), and modulus gradation between distinct anatomical regions of the thorax for asynchronous (honey bee, *Apis mellifera, A. mellifera*) and synchronous (hawkmoth, *Manduca sexta, M. sexta*) fliers. We further utilized finite element analysis (FEA) to evaluate the impact of modulus gradient on thorax stress concentrations.

## 2. Experimental Methods

### 2.1 Insect care and selection of regions of interest

*M. sexta* larvae were sourced from Josh’s Frogs (Owosso, MI). Larvae were kept in inverted 0.95 liter plastic insect rearing cups (4 larvae per cup) with gutter mesh used as a climbing matrix. Rearing cups were inspected and cleaned daily. The rearing room was maintained at an ambient temperature of 28 ± 2 °C and larvae were subjected to a 24:0 (L:D) photoperiod to prevent pupal diapause [16]. Larvae were fed Repashy Superfoods Superhorn Hornworm Gutload Diet from Repashy Ventures (Oceanside, CA). Once the dorsal aorta appeared (7-14 days), larvae were moved to a pupation chamber, a large plastic bin with a layer of damp organic potting soil and gutter screen to facilitate adult wing unfurling. Adults emerged within 14-30 days and were sacrificed within two days of emergence with ethyl acetate in a kill jar. *A. mellifera* specimens were collected from a pollinator garden in Bozeman, MT. After euthanization, most specimens were preserved in 70% ethanol except for *A. mellifera* used in histology which were preserved with 10% neutral buffered formalin overnight.

To determine whether thorax properties are associated with anatomy, we assigned regions to the thorax. The *A. mellifera* thorax was divided into 4 regions (**Figure 1B**): The scutum (B1), the scutellum (B2), the postscutellum (B3), and the posterior phragma (B4) [17]. The posterior phragma extends internally to provide an attachment site for the dorsolateral flight muscles. *M. sexta* was divided into 3 regions (**Figure 1C**): anterior scutum where the dorsal lateral muscles attach (M1), the dorsal scutum where the dorsal ventral muscles attach (M2), and the posterior phragma, the second site for dorsal lateral muscles attachment (M3).

**Figure 1.**
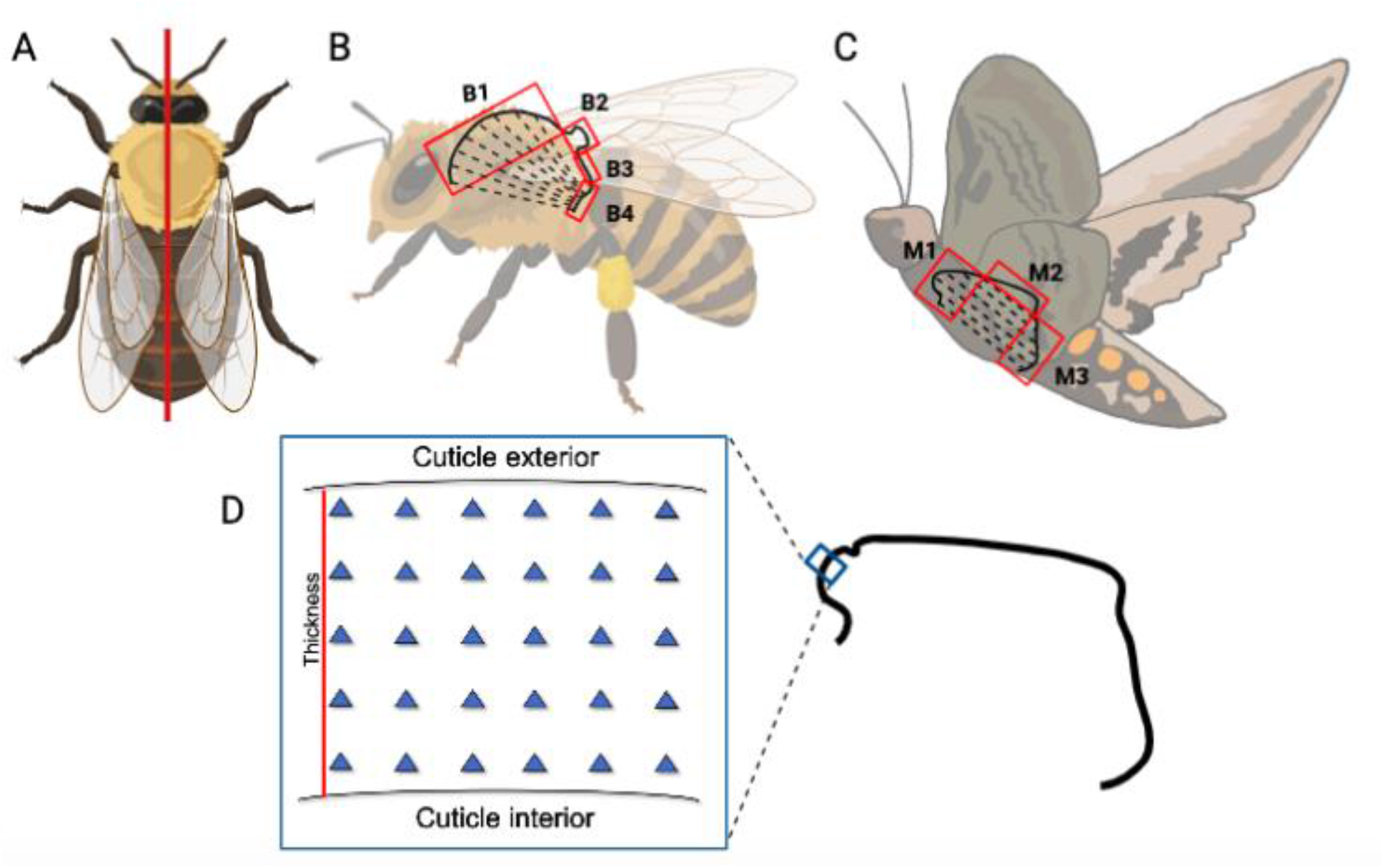
Approach for thoracic cuticle nanoindentation. (A) Insects were cross-sectioned in the sagittal plane to expose the dorsolateral muscle attachment sites and facilitate nanoindentation mapping of modulus gradation through the cuticle thickness. (B) The *A. mellifera* thorax was assigned functional regions B1-B4, where dashed lines show the dorsal lateral muscles. (C) *M. sexta* was assigned functional regions M1-M3. (D) The thickness-normalized position of each indent was calculated with reference to the cuticle interior (0 = interior, 1 = exterior).

### 2.2 Specimen preparation and histology

Histological staining was used to identify variation in thoracic cuticle structure. Formalin fixed *A. mellifera* specimens were dehydrated in a graded 70-100% ethanol series, embedded in paraffin, sectioned in the sagittal plane in 5 μm sections, and stained with hematoxylin and eosin (H&E)[18]. *M. sexta* specimens were stained using the same protocol but were not fixed with formalin. Fixation did not affect cuticle staining for either insect (**Supplemental Figure 1**) but improved the sectioning integrity for *A. mellifera*. Sectioned stained cuticles were imaged with a Nikon Eclipse E800 microscope using the infinity 2 color microscope camera. White balance was set to Red = 1.33 Blue = 1.00 and Green = 2.00. Regional cuticle thickness was measured using ImageJ 1.53a.

### 2.3 Specimen preparation and nanoindentation

Separate insects than studied for histology for both species were prepared for nanoindentation analyses. Insects were dehydrated via a graded ethanol series (70-100%; at least 3 days between each step), cleared with acetone, and embedded in polymethyl methacrylate (PMMA) at 35°C [19]. Samples were cross-sectioned in the sagittal plane, which was selected to expose the dorsolateral flight muscles and their cuticular attachment sites (**Figure 1A**). Cross-sectioning the cuticle facilitated mapping modulus gradation through the cuticle thickness while avoiding possible substrate effects that can occur from ‘top down’ indentation of thin structures of layers of different moduli [20,21]. The cut surface was polished with CarbiMet SiC 600 and 1000 grit abrasive paper lubricated with water, and then with progressively finer (9 to 0.05 μm) Ted Pella water-based alumina polishing suspension to a mirror finish. Samples were sonicated in water between polishing steps.

Nanoindentation (KLA-Tencor iMicro, Berkovich tip) was performed in load-control to a maximum load of 2 mN. The load function was 30s ramp, 60s hold, 10s unload. The 60s hold was sufficient to achieve dissipation of viscous energy, as confirmed using time-displacement plots. Non-overlapping nanoindentation arrays spanned the thorax thickness with 5 μm spacing between indents (**Figure 1B**). This spacing was chosen to maximize resolution of modulus gradient throughout the cuticle while maintaining approximately 3 times the contact radius between indents to minimize the impacts of the plastic zone between indents [22]. At least two non-overlapping arrays were placed in each thoracic region.

The reduced modulus (*E_r_*) was calculated using the Oliver-Pharr method (Equation 1) [23]. A second-order polynomial was fitted to the 95^th^ – 20^th^ percentile of the unloading portion of the load-displacement curve. Stiffness, *S*, was calculated as the slope of the tangent line to the start of the polynomial fit. The tip contact area (*A_c_*) was calibrated using fused silica and measured from contact depth.

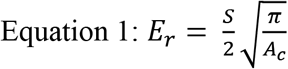

All indents were analyzed except for those that (1) were not on thorax (e.g., in plastic or on cuticle/plastic interface, determined through inspection with a 50x microscope), (2) did not demonstrate a smooth elastic-plastic transition, or (3) did not demonstrate a smooth loading-unloading curve. The position of each indent was calculated from the relative distance of the residual indent from the outer surface (0 = cuticle interior, 1 = cuticle exterior) (**Figure 1D**).

### 2.4 Confocal Scanning Laser Microscopy

Because cuticle can contain resilin and resilin stiffens when dehydrated [24,25], we sought to assess whether cuticle with higher moduli also had higher resilin content. Resilin is autofluorescent, with excitation of ~350-407 nm and emission of ~413-485 nm [26]. Following nanoindentation, samples were imaged for resilin fluorescence with a Leica TCS SP5 (excitation 405 nm, emission 420-480 nm) [27]. Imaging was performed using a 25x long working distance water immersion objective. Images were processed using Imaris 9.3.0.

### 2.5 Finite Element Analysis

We developed a simple 2D FEA model (ABAQUS CAE 2019) to evaluate the impact modulus gradients had on stress in deformed cuticle (**Figure 2**). The cuticle was treated as a rectangle of length 1000 μm long and height 30 μm. The height was assigned as 30 μm because this value is between the mean cuticle thickness measurements for *A. mellifera* (17 μm) and *M. sexta* (45 μm). The cuticle was discretized into 1660 linear quadrilateral S4R elements, which was sufficient for convergence of maximum stresses in all cases. For simplicity, only half the cuticle structure was modeled, where the centerline was constrained to only allow deformation along the y-axis. The lower edge was constrained to only deform along the x-axis. A small deformation of 5% of the cuticle thickness was prescribed in the negative y-direction at the top edge of the cuticle section. This deformation magnitude was chosen to ensure that the model remained in the geometrically linear regime. The cuticle was deformed 5% at 0 μm and the deformation linearly decreased so that the cuticle end (500 μm) was not deformed. To our knowledge, the thoracic cuticle deformation (which is not identical to bulk thorax deformation) during flight has not been measured in any insect. Stress depends on material properties as well as thorax geometry. By choosing modulus and cuticle thickness values that are representative of experimental values for *A. mellifera* and *M. sexta* we gain a qualitative understanding of how modulus gradients affect stress accumulation in insect cuticle.

**Figure 2.**
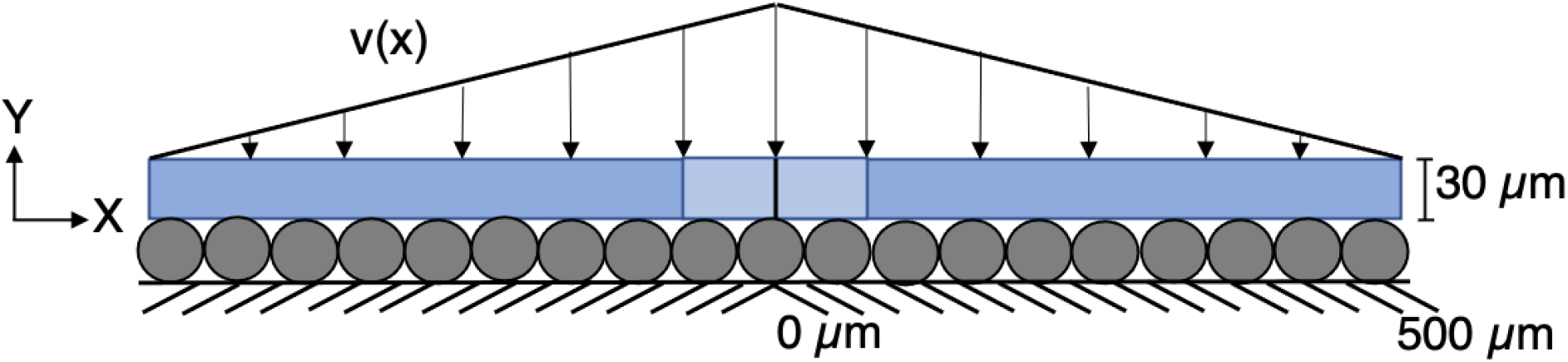
FEA schematic of a 2D insect cuticle. v(x) denotes a prescribed displacement field changing in the x-direction. We focused our discussion on middle 5% of the cuticle where the highest stress occurred (light blue).

We assigned one of two elastic moduli distributions. The first was a uniform distribution, and the second was a 2 GPa linear gradient in the y-direction. We considered a positive modulus gradient to represent forces that may be applied from the exterior such as during predation or burrowing. We also considered a model with a negative modulus gradient to represent forces that may be acting from the interior such as flight muscles. In all cases the cuticle material was assumed linear-elastic and isotropic. We focused on the area with the largest stress values by zooming in on the 5% of the cuticle with the largest imposed displacement. Results were mirrored to show the effect of deformation in both directions.

### 2.6 Statistics

Between 2 and 7 nanoindentation arrays were collected for each region. To avoid oversampling some regions, two arrays were randomly selected from each region for each specimen for analysis. We sought to test the hypotheses that (1) mean modulus and (2) modulus gradation through the cuticle thickness vary by region. To test our first hypothesis, we generated a mixed model ANOVA with the random effects of specimen and array and the fixed effect of region. To test our second hypothesis, we used a linear mixed model with random effects of specimen and array, fixed effect of region, and a covariate of indent relative position. Relative positions were assigned within the thorax thickness, where 0 indicates the cuticle interior and 1 indicates the cuticle exterior. Modulus was natural log transformed, when necessary, to satisfy ANOVA assumptions of residual normality and homoscedasticity. All statistical tests were performed using Minitab v. 19 2020 2.0. The threshold for significance was set *a priori* at p < 0.05 for all tests.

## 3. Results

### 3.1 Histology

H&E staining showed that the thorax composition is heterogeneous between regions for *A. mellifera* and *M. sexta* (**Figure 3**). All *A. mellifera* regions had distinct exocuticle and endocuticle (noted by the stark difference in pink color) except for region B1, where these layers were not distinct. *M. sexta* had distinct endocuticle and exocuticle separation in M1 and M2 but not M3. *M. sexta* epicuticle is visible as a thin dark line on the cuticle exterior, which is not seen in *A. mellifera*. The epicuticle may not be visible with the dark staining of the exocuticle. *A. mellifera* exocuticle showed dark exocuticle staining for all regions. A similar dark staining is seen for *Drosophila melanogaster* abdomen cuticle[28] and may be associated with highly sclerotized cuticle[29,30].

**Figure 3.**
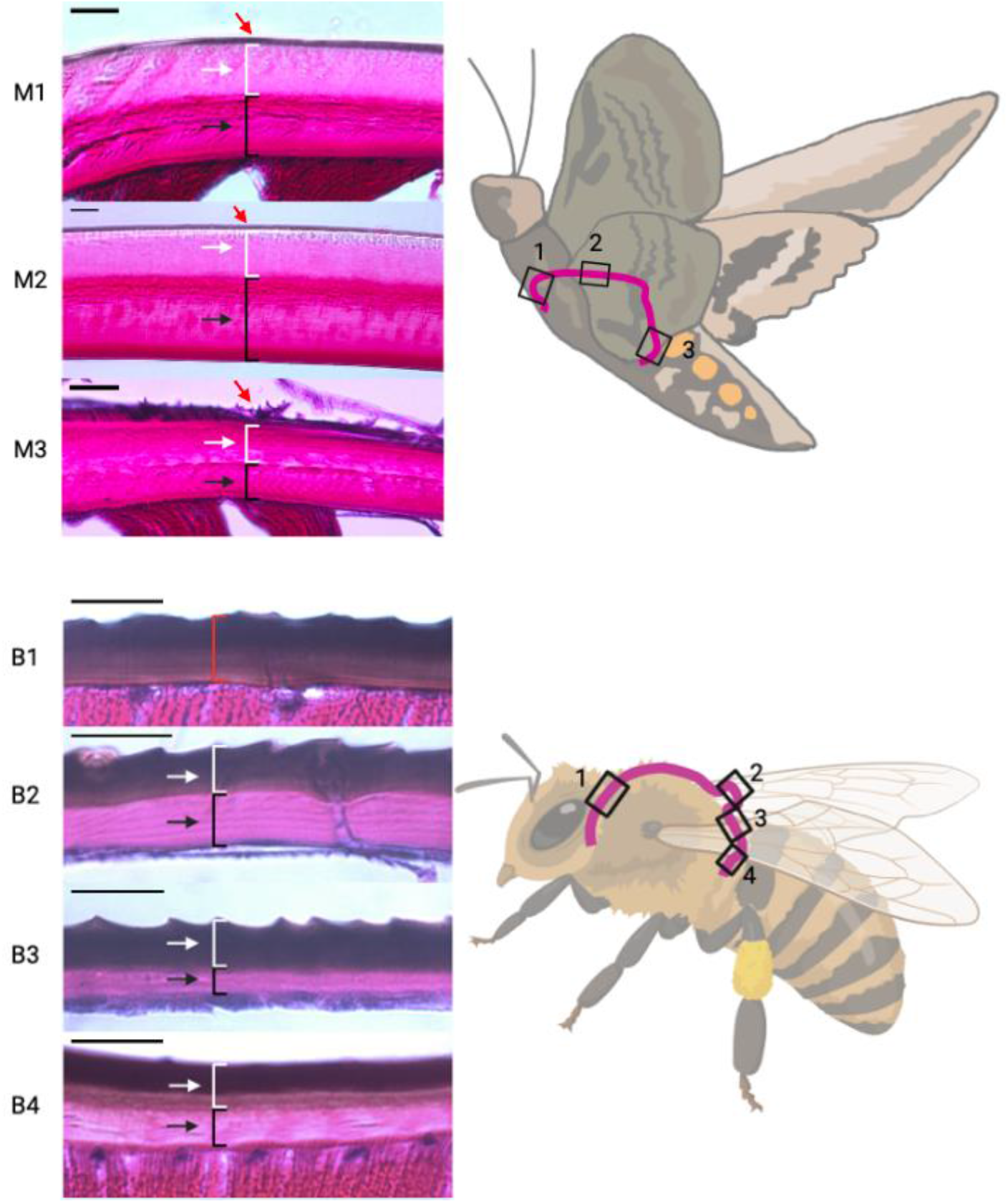
Thorax structure from H&E stained sections differs by insect and region. For all images, the cuticle interior is on the bottom. Endocuticle (black arrows) and exocuticle (white arrows) thicknesses vary between thorax regions for *A. mellifera* and *M. sexta*. An epicuticle (red arrows) was only observed for *M. sexta*. *A. mellifera* and *M. sexta* cuticle samples were imaged at 600x and 200x, respectively. Scale bars are 20 μm.

For each insect, the thorax composition varied by region (Table 1). For *A. mellifera*, the cuticle thickness increased as B3 (15.5 μm), B1 (16.8 μm), B4 (17.4 μm), and B3 (19.9 μm). B3 had a much thicker exocuticle than the other regions. We note that cuticle layering was more difficult to measure in B1, where layers were not distinct. Epicuticle was not detected for *A. mellifera*. For *M. sexta*, M1(49 μm) and M2 (52 μm) had similar thicknesses while M3 was only 33 μm, 40-45% thinner than the other regions. The epicuticle in *M. sexta* ranged from 4-9%. M1 had the thickest endocuticle at 55%.

**Table 1.** Cuticle layer composition by region for *M. sexta* and *A. mellifera*.

| Cuticle Layer % | M1 | M2 | M3 | B1 | B2 | B3 | B4 |
| --- | --- | --- | --- | --- | --- | --- | --- |
| Epicuticle | 6 | 4 | 9 | - | - | - | - |
| Exocuticle | 41 | 54 | 46 | - | 45 | 64 | 41 |
| Endocuticle | 53 | 42 | 45 | - | 55 | 36 | 59 |

### 3.2 Nanoindentation

We used nanoindentation to evaluate the dependence of mean cuticle modulus and modulus gradation on thorax region. The mean modulus did not differ between region for either *A. mellifera* or *M. sexta* (p > 0.05 for both) and did not differ between *A. mellifera* (7.00 ± 1.29 GPa) and *M. sexta* (6.98 ± 1.02 GPa) (p > 0.05).

Nanoindentation modulus was graded with respect to cuticle position for *A. mellifera* (**Figure 4**). This gradation was approximately linear, increasing from the cuticle interior (relative position = 0) to the exterior (relative position = 1) (**Supplemental Figure 2)**. We used mixed model ANOVA to evaluate whether these gradations differed between regions for *A. mellifera*. There was a significant (p < 0.001) interaction between region and relative position, indicating that the linear gradient of modulus (i.e., slope) differs between regions. Equation 2 describes the marginal fits for Er (GPa) by region for *A. mellifera* from mixed model ANOVA.

**Figure 4.**
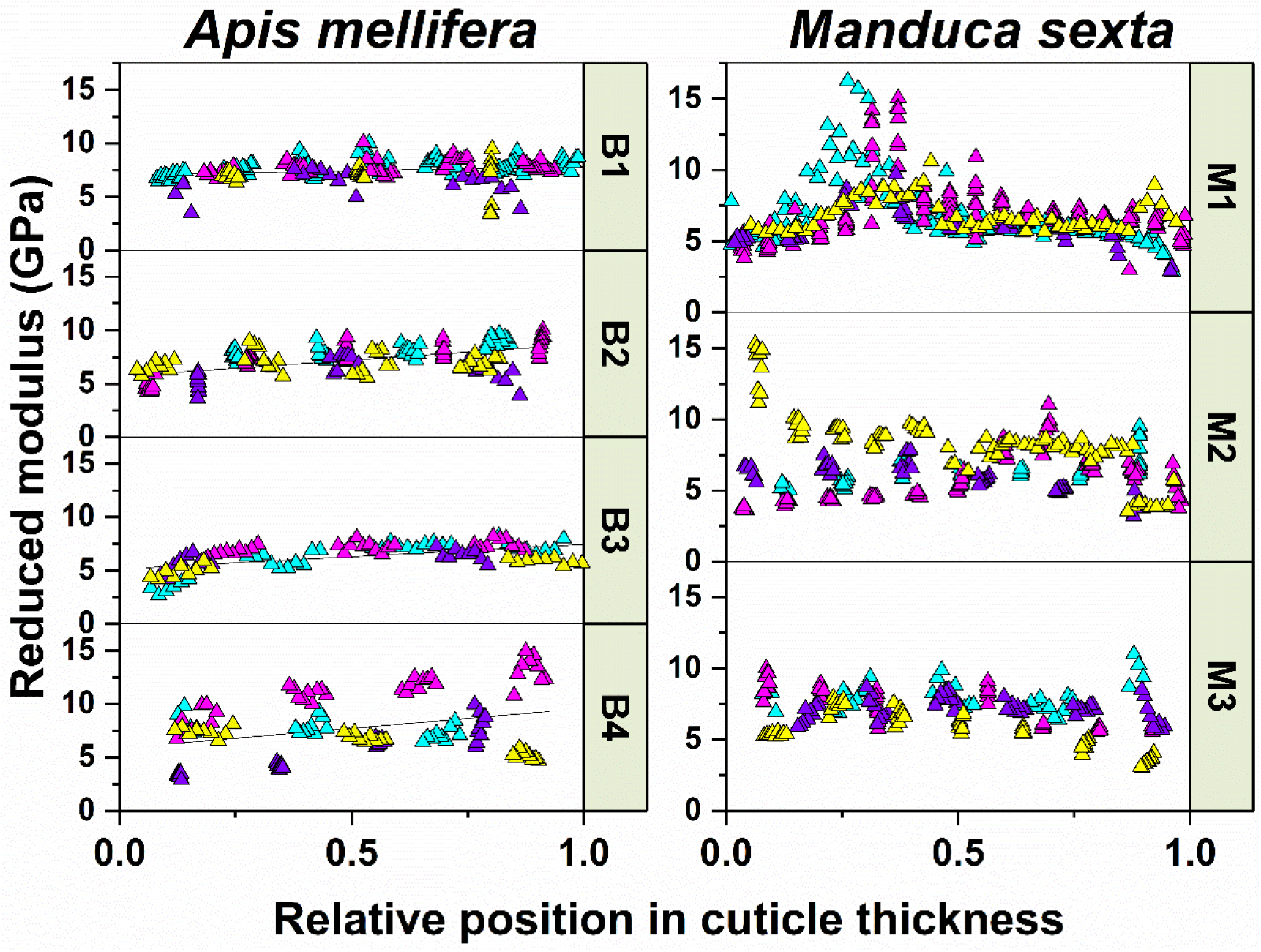
Modulus gradation through the cuticle thickness where 0 indicates cuticle interior and 1 indicates exterior. Data are those analyzed for *A. mellifera* and *M. sexta*. Each color represents all indents from one array per region. *A. mellifera* modulus gradients were well-represented by linear fits for all regions. Modulus did not show consistent gradation for *M. sexta* in any region.

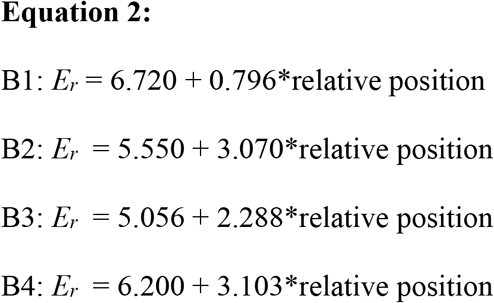

The slope in B1 was close to 0 and was much less (97-118%) than in B2-B4. While a single estimate of slope represented most of *A. mellifera* regions well (**Figure 4**), B4 was more variable. Half of the arrays in this region had a steep positive slope and half had a slight negative slope which resulted in a mean positive slope. Modulus gradients for *M. sexta* were variable in both direction and shape with no single approximation appropriate for the regional representation of the data.

### 3.3 Confocal laser scanning microscopy

Because dehydrated resilin may stiffen the cuticle [24,25], we employed CLSM to determine whether modulus gradients correspond with resilin distribution. Resilin identified from blue autofluorescence was present in the cuticle of both insects but did not correspond with the locations of higher moduli (**Figure 5**). For *A. mellifera*, the presence and location of resilin was inconsistent between specimens. B1 did not have discernable resilin (images demonstrate only autofluorescence of muscle), while regions B2-B4 showed resilin for one specimen but not the other. Resilin was present for both specimens at the scutellum hinge (red arrow), as expected given the high level of articulation of this structure. *M. sexta* had resilin throughout the thorax cuticle but composition varied by region. In particular, M2 lacked resilin on the interior portion of the cuticle.

**Figure 5.**
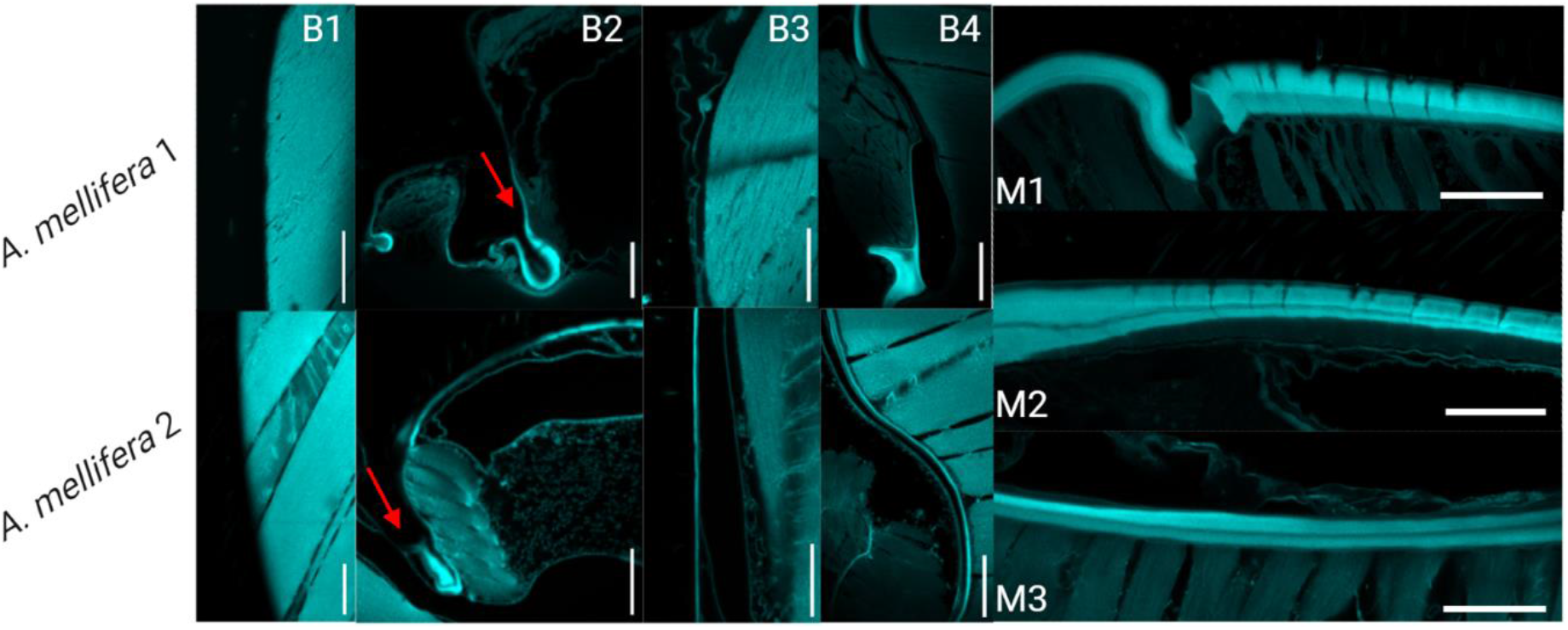
Confocal laser scanning microscopy showed autofluorescent resilin in the insect cuticle. In *A. mellifera*, resilin content varied between the two specimens (top and bottom). B1 lacked resilin while B2-B4 had resilin in one specimen but not the other (cuticle interior on right). The scutellum was resilin rich for both specimens (red arrows). *M. sexta* had resilin throughout the cuticle but distribution varied by region (cuticle interior on bottom). Images were taken at 25x using a 405 nm laser collecting emission from 420-480 nm. Scale bars are 100 μm.

### 3.4 Finite element analysis

FEA was used to investigate how experimentally-observed modulus gradients influenced stress distributions in the deformed cuticle. A homogenous cuticle was first modeled to provide a baseline calculation of stress distribution. The cuticle was assigned a modulus of 7 GPa from the mean experimental data. A second model considered elastic moduli that varied linearly from 6 to 8 GPa through the cuticle thickness. A slope of 2 GPa increase across the cuticle thickness was chosen based on an average estimate of slope for *A. mellifera*. The slope was applied in the positive y-direction and, in a separate model, the negative y-direction based on the potential for forces from flight muscles or from external sources to be acting on the cuticle.

The linear modulus gradient influenced the maximum stress, but the stress distribution was insensitive to modulus gradient or direction (**Figure 6**). When the deformation was applied to the high modulus edge, the linearly-graded cuticle experienced a maximum stress 15.3% less than the homogenous cuticle. On the other hand, when the deformation was applied to the low modulus edge, the linearly-graded cuticle experienced a maximum stress 13.3% greater than that experienced by the homogeneous cuticle. This complexifies the stress reduction in the thorax cuticle. Based on our simplified FEA model, a linearly distributed modulus cannot optimally reduce stress for both interior deformation due to flight muscles and exterior deformation from external forces.

**Figure 6.**
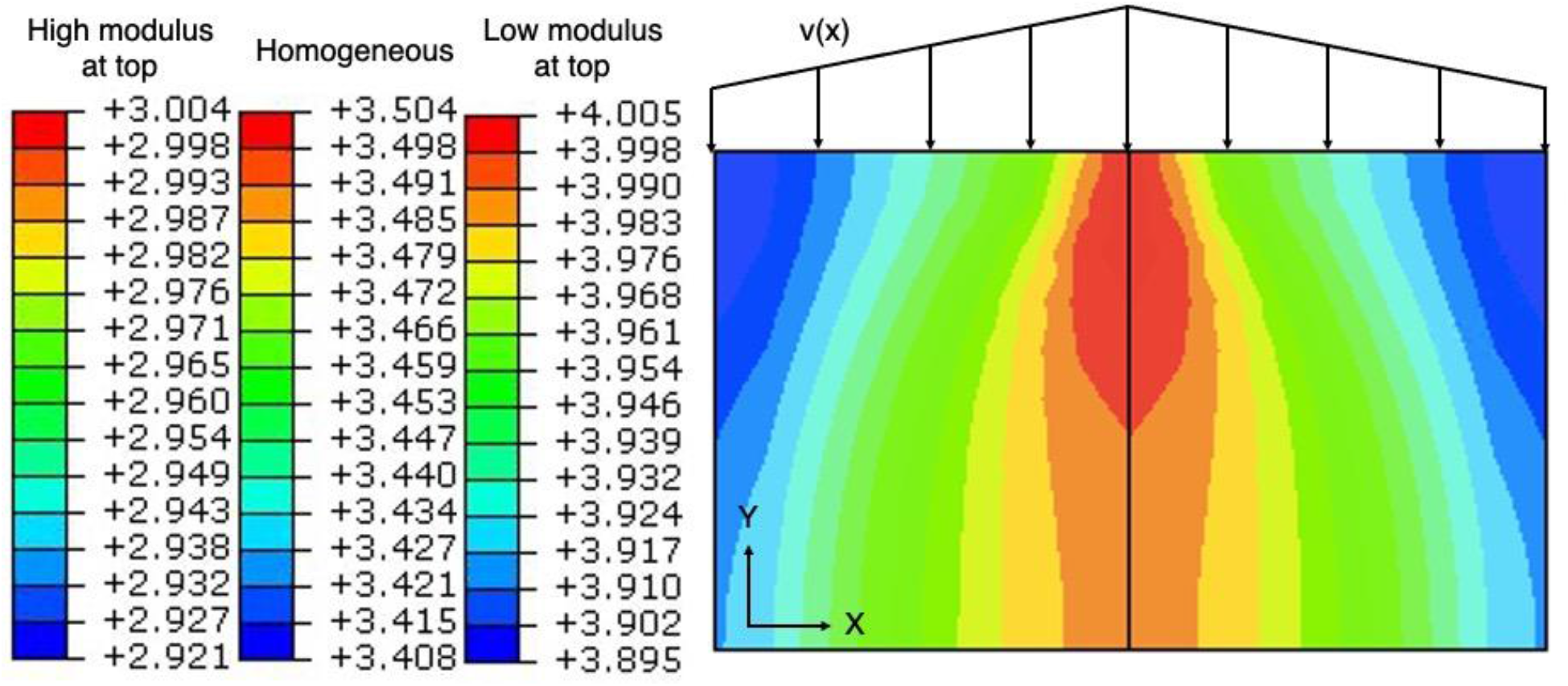
Stress distribution (10^8^ Pa) from a 5% deformation of 2D thoracic cuticle models. The lowest stress accumulation occurred when the higher modulus edge of the linear distribution was deformed.

## 4. Discussion

The thorax is essential for insect flight, yet significant questions remain about its exceptional mechanical properties. The thorax must be flexible enough to articulate the wing but must also protect against external forces and resist plastic deformation under the high magnitude forces produced by the flight muscles. Furthermore, the thorax stores and releases elastic energy over a large number of oscillations (e.g., 10^7^ oscillations for *A. mellifera* [31]) without fatigue. Modulus gradients are observed in other biological materials (e.g., enamel, bamboo, nacre) where they help maintain a balance of strength, resilience, and toughness and minimize stress concentrations in complex geometrical structures [32–35]. The purposes of this study were to determine whether insect thorax has graded moduli through the cuticle thickness, whether these gradients depend on thorax region and species, and whether the modulus gradients impact thorax stress concentrations.

We performed nanoindentation on ethanol-dehydrated, plastic-embedded samples of thorax cuticle. The mean moduli for thorax cuticle in both *A. mellifera* and *M. sexta* were in good agreement with values reported for different insect structures. Our measurement of ~7 GPa for dehydrated thorax cuticle for *A. mellifera* and *M. sexta* is within the 4.8 to 9.9 GPa reported for the beetle gula [6–8], elytra [9,36], and infrared sensillum [7], and the locust sternum [13] and tibia [12]. These are among the first nanoindentation evaluations for the flying insect thorax cuticle, and specifically within the *Hymenoptera* and *Lepidoptera* taxa [37,38]. Studies comparing nanoindentation results from dried and fresh cuticle of the beetle gula [6,8], locust sternum [13], and termite mandibles [39] showed modulus increases of 10% to upwards of 500% with dehydration. Our thorax moduli for *M. sexta* were ~25% greater than for modulus values of the fresh thorax reported for the same species [38]. This level of stiffening from dehydration is on par with that observed in bone embedded in PMMA [19]. We find this to be an acceptable tradeoff for high spatial resolution testing which is difficult to achieve in fresh samples because of higher surface roughness. The high modulus values are unlikely to be driven by dried resilin content. Our CLSM imaging shows that there are regions of cuticle with more resilin, but that those regions are not consistent with regions that exhibited increased modulus.

Modulus varied throughout the thorax thickness, but the direction and steepness of this gradient depended on insect and region. For two of the four regions, *A. mellifera* had steep, positive, linear, modulus gradients from cuticle interior to exterior. The positive modulus gradient in *A. mellifera* thorax agrees with gradients found in non-thorax cuticle for some other insects. Previous work demonstrates that the modulus increases from the cuticle interior to exterior for the gula [6–8], infrared sensillum [7], and elytra [9], and tarsal setae[10] of various beetles, the grasshopper mandible[11], and the locust tibia[12] and sternum[13]. Increased modulus is observed with sclerotization [40,41], including for some of these insect structures [7,12]. For *A. mellifera*, the dark staining at the thorax exterior from histological sections indicates that higher modulus is also likely to be influenced by sclerotization. However, sclerotization cannot be the only contributor to modulus gradation, since dark banding was seen on the exterior of all *A. mellifera* histological sections but did not always align with the higher modulus seen at the exterior of the same regions. The apparent lack of sclerotization in the cuticle for *M. sexta* may partially explain the differences in modulus gradation between *M. sexta* and *A. mellifera* found in this study. The differences between *A. mellifera* and *M. sexta* thorax cuticle may also be adaptations for behaviors separate from or in conjuncture with flight. For example, most cases of increasing modulus gradient, as seen in *A. mellifera*, are found for cuticle optimized for protection against predation or during burrowing [9,15,42]. Although *A. mellifera* do not burrow, most *Hymenoptera* do (e.g., genera *Xylocopa, Andrenida* [43]). Thus, the protective cuticle may be residual in the Hymenoptera lineage to adapt for burrowing.

We used FEA to assess the impact of the modulus gradient found in *A. mellifera* on cuticle stress concentration. Lower stress accumulation is essential for avoiding permanent deformations that could impact thorax performance. The cuticle accumulated the least stress when its modulus was distributed linearly through the thickness, and it was compressed from the highest modulus side. For *A. mellifera*, this type of compression would likely result from external forces, which fits with our hypothesis that the cuticle gradient is optimized for protection. While simplified, these models illustrate that graded moduli distribution can reduce stress accumulation in some loading contexts. However, further work needs to be done to understand how bulk thorax geometry coupled with cuticle material variation influence peak stresses and energy storage during flight.

The thorax likely uses several mechanisms to increase energy return during flight. For both synchronous (e.g., *M. sexta*) and asynchronous (e.g., *A. mellifera*) fliers, large power requirements, and low muscular efficiency (<10%) necessitate energy storage in the thorax [44]. Passive, asynchronous muscle is stiffer and behaves more spring-like relative to synchronous muscle [45]. This passive stiffness may be one way that asynchronous muscle stores elastic energy. In *M. sexta*, mechanical coupling of the scutum and the wing hinges is believed to increase energy return by decoupling the area where the most power is lost (wing hinges) and where the most energy is returned (scutum) [46,47]. Material gradients may also play a role in this elastic energy storage. Modulus gradients can increase energy return in biomaterials and bioinspired polymers by increasing stretching [34,35]. Further research is needed to identify how these properties or others adapt the thorax cuticle for energy storage in different flight lineages.

While this study has elucidated the material properties and structure of flight insect thorax cuticle, there are several limitations to our approach. First, modulus is affected by many factors, including material composition and organization [12,13,48].We identified but did not quantify resilin or sclerotization, which is not possible using CLSM. Second, dehydrating the cuticle was necessary to achieve high spatial resolution for nanoindentation. However, dehydration stiffens the cuticle and may affect material gradients [10,13].

### 4.1 Conclusions

We demonstrate that the flying insect thorax is heterogeneous with respect to structure, material composition, and modulus gradient. In *A. mellifera*, a clear linear modulus gradation is observed in two of the four anatomic regions of interest within the thorax. Thorax assessment at one location may not adequately capture variation occurring within and between flying insects. While the linear gradient of modulus with thorax thickness may contribute to decreased stress concentration, further work is needed to determine the relative contributions of cuticle structure, material composition, and modulus gradient to the efficiency of different forms of flight.

## Acknowledgements

We would like to thank Robert KD Peterson and Miles Maxcer for helping raise insects, and Ghazal Vahidi for assistance in sample preparation for nanoindentation.

## Author contributions

<u>Study design</u>: CC, CY, MJ, CH; <u>experimental data collection</u>: CC, <u>model development</u>: CC and MJ, <u>care and raising of insects</u>: CY, <u>data analysis</u>: CC and CH, <u>funding</u>: MJ, <u>drafting manuscript</u>: CC, MJ, CH, <u>final editing and approval of manuscript</u>: CC, CY, MJ, CH.

## Funding statement

This research was partially supported the National Science Foundation under awards No. CMMI-1942810 to MJ. Any opinions, findings, and conclusions or recommendations expressed in this material are those of the author(s) and do not necessarily reflect the views of the National Science Foundation.

## Supplemental materials

**S1.**
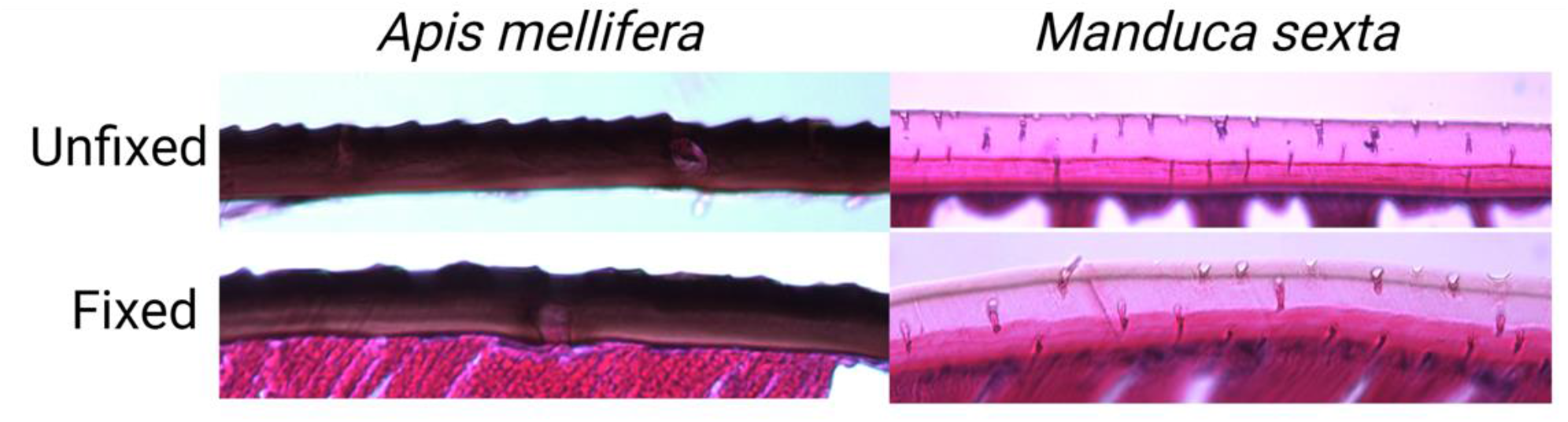
Using fixed or unfixed insect samples did not change the staining. But fixation was necessary for *A. mellifera* because the cuticle tended to fall apart when unfixed. *A. mellifera* images were taken at with a 50x extra-long working distance lens (500x total magnification) and *M. sexta* images were taken at 200x.

**S2.**
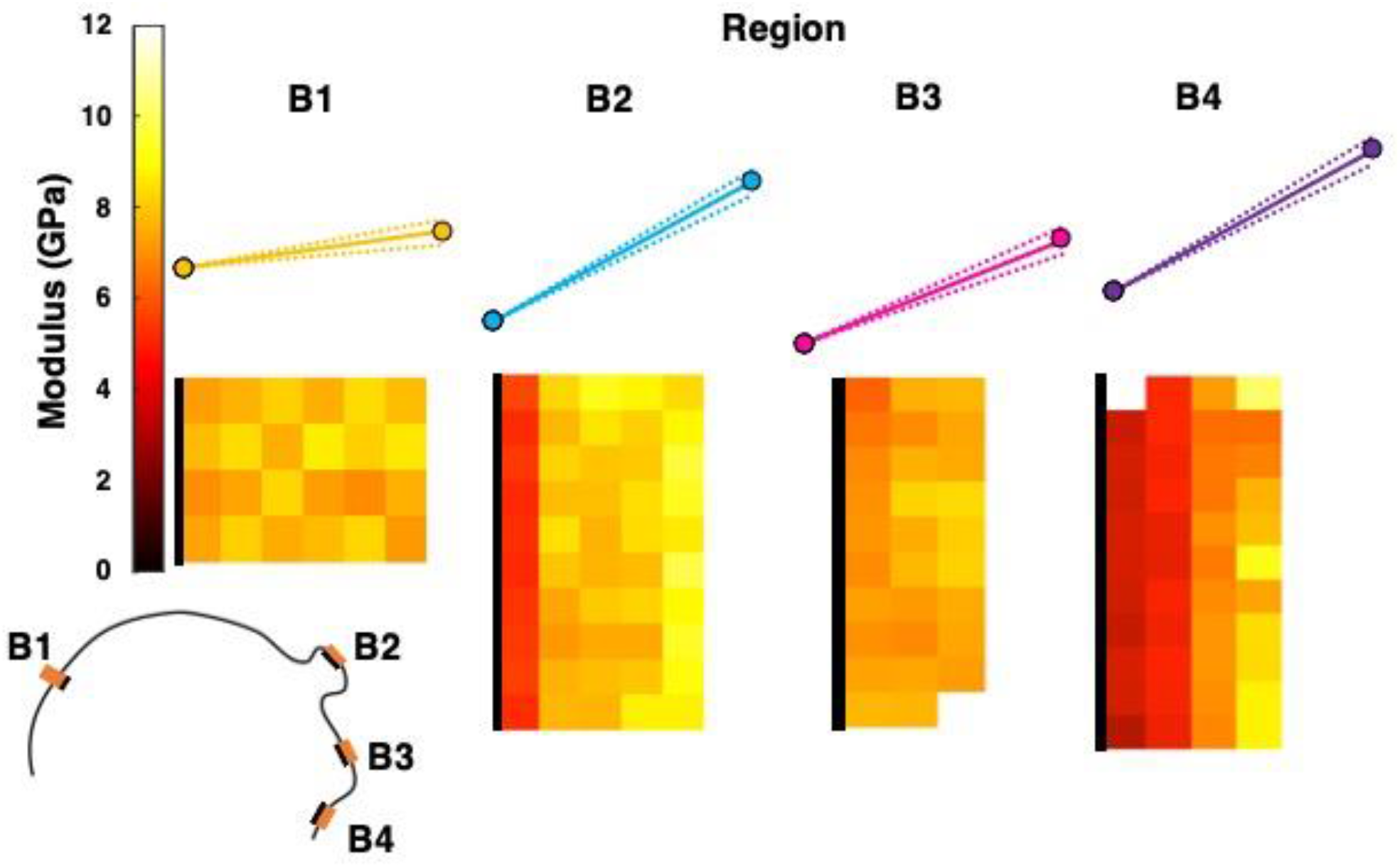
Plotted mixed model ANOVA equations with representative heatmaps for each region demonstrating the change in modulus through the thickness of the cuticle. Dashed lines represent standard error for slope from mixed model ANOVA. The black line on heatmaps indicates cuticle interior. While a single gradient represented most regions well, it did not for B4. In this region, half of the arrays had a steep positive gradient and half had a slight negative gradient which resulted in a mean positive slope.

## Notes

### Competing Interest Statement

The authors have declared no competing interest.

